# Exocyst subcomplex functions in autophagosome biogenesis by regulating Atg9 trafficking

**DOI:** 10.1101/306969

**Authors:** Sunaina Singh, Sarika Chinchwadkar, Amol Aher, Saravanan Matheshwaran, Ravi Manjithaya

## Abstract

During autophagy, double membrane vesicles called autophagosomes capture and degrade the intracellular cargo. The *de novo* formation of autophagosomes requires several vesicle transport and membrane fusion events which are not completely understood. We studied the involvement of Exocyst- an octameric tethering complex, which has a primary function in tethering post-Golgi secretory vesicles to plasma membrane, in autophagy. Our findings indicate not all subunits of exocyst are involved in selective and general autophagy. We show that in the absence of autophagy specific subunits, autophagy arrest is accompanied by accumulation of incomplete autophagosome-like structures. In these mutants, impaired Atg9 trafficking leads to decreased delivery of membrane to the site of autophagosome biogenesis thereby impeding the elongation and completion of the autophagosomes. The subunits of exocyst which are dispensable for autophagic function do not associate with the autophagy specific subcomplex of exocyst.

## Introduction

Autophagy is an intracellular catabolic process involving capture of cytosolic cargo by double membrane vesicular structures called autophagosomes. These autophagosomes then fuse with lysosomes (vacuoles in yeast) leading to degradation of the cargo (1). Studies by various groups have led to the identification of more than 40 ATG proteins (**A**utophagy Related **G**enes) and several accessory components. The molecular mechanisms of how these proteins function in the process of autophagy have been deciphered to a substantial extent (2). Autophagosome biogenesis is a complex process which begins with the assembly of autophagy initiation complex (Atg1 complex) at PAS (Pre-autophagosomal structure, a perivacuolar autophagosome biogenesis site in yeast) followed by activation of VPS34 complex at this site to produce PI3P locally thereby leading to the recruitment of other core autophagy proteins and nucleation of precursor autophagosome membrane. This nascent structure known as the phagophore further expands into double membrane vesicle by addition of membrane derived from various sources mediated by Atg9 vesicles (3, 4).

Atg9 is an integral membrane protein which appears as multiple puncta in cytoplasm (5). These puncta represent the peripheral pool of Atg9 containing vesicles which deliver membrane to PAS allowing autophagosome expansion. These Atg9 vesicles are known to be derived from various membrane sources including Golgi associated secretory pathway (6–11).

Various secretory pathway proteins were shown to be important for the process of autophagosome biogenesis. The early secretory pathway components consisting of multisubunit protein complexes such as COPII machinery, COG, GARP,TRAPPIII, TRAPP IV, late secretory components such as Sec2, Sec4 and various SNARE proteins such as Sso1, Sso2, Sec9,Tlg2, Ufe1, etc. were shown to be important for autophagosome biogenesis (12–16). Among these, several tethering complexes like COG, TRAPIII and exocyst have been associated with autophagy. While TRAPIII function has been elucidated to an extent, the involvement of other two complexes is not clear (17–21). Exocyst is a conserved multisubunit protein complex which functions during exocytosis (22, 23). This octameric complex consists of Sec3, Sec5, Sec6, Sec8, Sec10, Sec15, Exo84 and Exo70 subunits (24–26). It helps in tethering post-Golgi secretory vesicles to the plasma membrane (27). In yeast, the exocyst subunit Sec3 serves as a landmark for the assembly of exocyst complex at the plasma membrane (28). This assembly is also regulated by Sec4, a Rab protein present on the secretory vesicle (29, 30). In mammalian cells, Moskalenko and colleagues suggested that the exocyst complex exists as two subcomplexes, one forms a targeting patch at the plasma membrane and the other helps in directing the secretory vesicles to the location marked by targeting patch. This coalition of both the subcomplexes is brought about by Ral-GTPases (31).This phenomenon of Ral mediated activation of exocyst complex was also recounted where the activation of Sec5subunit led to autophagy inhibition while activation of Exo84subunit led to induction of autophagy by assembly of initiation complex (32). In yeast, Rho GTPases are known to mediate the assembly of the exocyst complex (33). Some exocyst subunits are required for autophagy which underscores the overlap between the secretory pathway components and autophagy (16). It is suggested that the determinants of the membrane flow in the secretory pathway also contribute to autophagosome biogenesis (12). As exocyst is important for secretory function, it may play a similar role in autophagy where vesicular based membrane flow is critical. A study suggested that tethering function of exocyst is responsible for SNARE pairing in the secretory pathway (3, 4) and this idea was extended to exocyst function in autophagy (16).However, the exact role of exocyst in yeast during autophagy is not well understood. Here, we report that several but not all subunits of exocyst are essential for autophagy. Some of the participating subunits affect Atg9 trafficking suggesting a role for this subcomplex in autophagosome biogenesis by contributing to the membrane flow. Our biochemical investigations further provide insights into the function of a distinct subcomplex of exocyst as during autophagosome biogenesis.

## Results

### Exocyst complex is involved in selective and general autophagy

We screened a subset of temperature sensitive (Ts) *Saccharomyces cerevisiae* mutants defective in vesicular trafficking for their ability to perform selective autophagy of peroxisomes-pexophagy. We measured pexophagy in these Ts mutants by employing a previously established immunoblotting assay in which accumulation of free GFP with concomitant decrease of fused protein Pot1-GFP is indicative of pexophagy (35). The cells expressing Pot1-GFP were grown in rich medium (YPD) and actively growing cells were then incubated overnight in fatty acid rich medium (Oleate) to allow build-up of peroxisomes. Pexophagy was induced by moving cells into starvation medium (SD-N). Among the mutants which showed a block in pexophagy, there were several mutants of the exocyst complex. We observed that the temperature sensitive mutants of exocyst complex *sec3-2, sec5-24, sec6-4, sec8-6, sec8-9, sec10-2* and *exo84-102* showed accumulation of free GFP at permissive temperature (PT, 25°C) but failed to do so at non-permissive temperature (NPT, 37°C) indicating a block in pexophagy (Fig. 1A, Fig. S1A). However, we also observed that not all mutants showed this block. At NPT, while *sec15-1* showed reduced levels of pexophagy, *exo70-38* was not affected (Fig. 1A).

**Figure 1:**
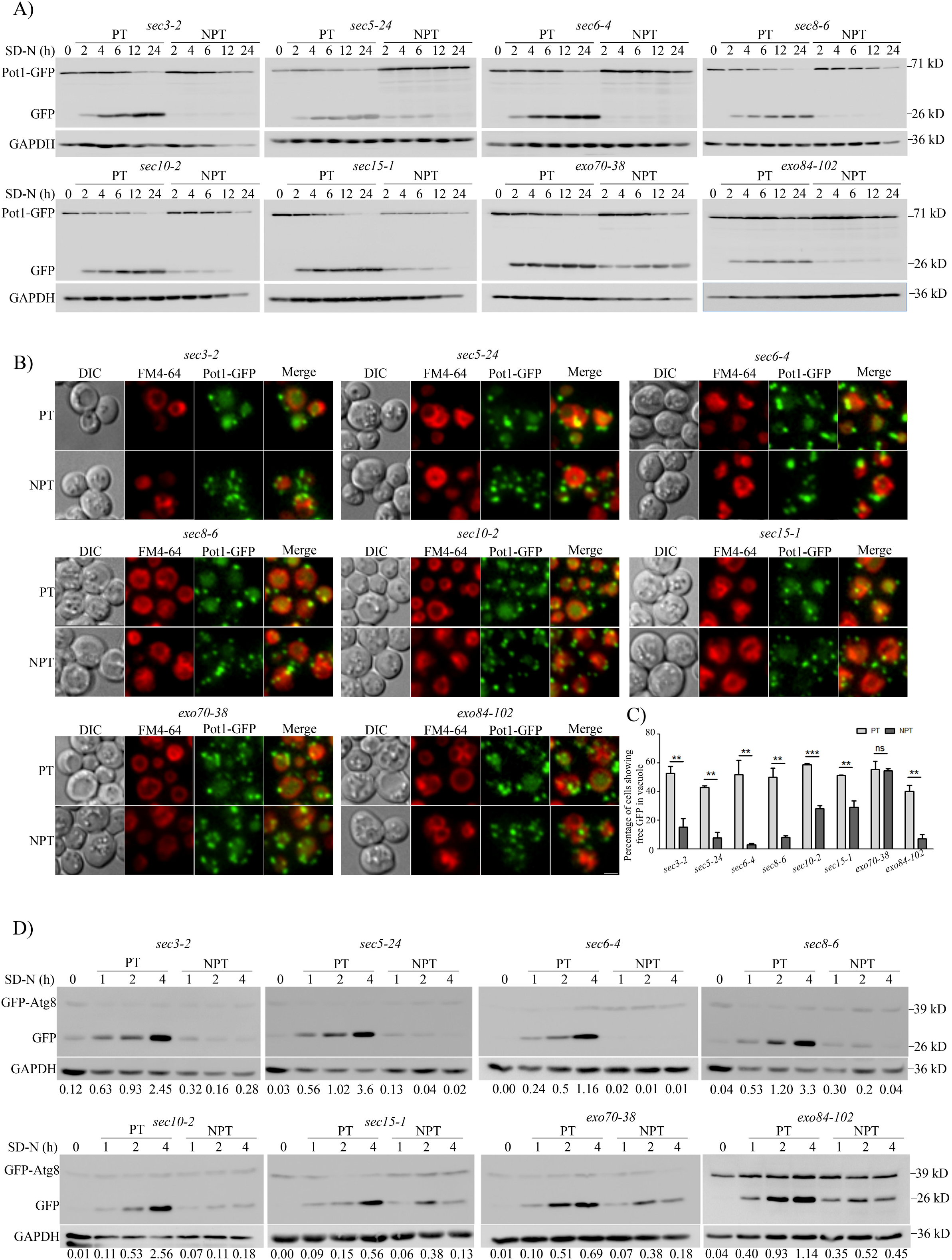
Subset of exocyst complex mutants are defective in selective and general autophagy. A) Temperature sensitive exocyst mutants expressing Pot1-GFPwere grown in oleate medium to induce peroxisomes. They were subsequently transferred to nitrogen starvation medium (SD-N) to induce pexophagy under permissive (PT) or non-permissive temperatures (NPT). Samples were collected at indicated time points, processed and subjected to immunoblotting analysis. B) Cells were treated as in A and were imaged 4 hours in starvation medium using fluorescence microscopy. Peroxisomes appear green due to the presence of Pot1-GFP. Vacuoles were labelled with FM4-64 dye. Images were deconvolved by nearest neighbour algorithm using softWoRx software (GE Healthcare) and maximum intensity projection images are shown. Scale bar: 2μm. C) Quantitation of pexophagy from images obtained in B. About 150-200 cells for each experiment were counted for the presence of GFP in the vacuole and represented as percentage of total cells scored. The bar diagram shows mean of three independent experiments with standard error. Statistical significance was analysed by Student’s unpaired t-test. ns-nonsignificant, **, *P*<0.01, ***, *P*<0.001. D) GFP-Atg8 processing assay for general autophagy. Cells expressing GFP-Atg8 were starved in SD-N medium at PT or NPT. Samples were collected at indicated time points, processed and analysed by immunoblotting. Numbers indicate ratio of intensity of free GFP/ GAPDH.

To further validate these results, we performed fluorescence microscopy based pexophagy assay with these mutants expressing Pot1-GFP. When grown in oleate medium, peroxisomes appeared as green punctate structures in the cells. These cultures were then resuspended in starvation medium and were incubated at permissive and non-permissive temperatures. After 4 hours of starvation at permissive temperatures, all mutants showed accumulation of free

GFP inside the vacuoles labelled in red with FM4-64 (Fig. 1B). In agreement with our western blot data, we observed that there was no free GFP in the vacuole of all the mutants which showed a pexophagy block at non-permissive temperature (Figs. 1B, 1C, S1B and S1C).

Next, we wanted to check whether this defect is specific to a selective autophagy process like pexophagy or extended to general autophagy as well. Cells expressing GFP-Atg8, an autophagy marker were grown in selection medium (SD-Ura), transferred to starvation medium and incubated at permissive and non-permissive conditions. Immunoblotting studies revealed accumulation of free GFP over the course of starvation at permissive temperatures while no such accumulation was observed at non-permissive temperatures in most of the mutants (Fig. 1D and S1D). Similar to the pexophagy results, we observed no defect in general autophagy in *sec15-1* and *exo70-38* mutants. Surprisingly, we noticed that general autophagy was not blocked in mutant *exo84-102.* Whether Exo84 is important for only pexophagy or even for general autophagy still needs to be addressed.

### Mutants of exocyst complex accumulate immature autophagosomes

To investigate the stage of autophagy in which the mutants were blocked, we used fluorescence microscopy to observe localization of GFP-Atg8 puncta. A single perivacuolar punctum of Atg8 marks the site of autophagosome biogenesis (PAS). During starvation conditions where autophagy is prevalent, autophagosomes formed at PAS are expected to immediately fuse with the vacuole. Atg8, which is also a marker for autophagosome membrane, gets degraded along with the autophagosomal cargo in the vacuole.

Along with wild type cells, we studied two mutants which at non-permissive temperature showed complete autophagy defect (*sec3-2, sec8-6*) and one with partial defect (*sec15-1*). Interestingly, *sec3-2* and *sec8-6* both showed an increased number of Atg8 puncta in the cells but no diffused GFP in vacuoles indicating that possibly autophagosomal structures were getting accumulated. In addition, these cells did not display diffused GFP in the vacuole (Fig. 2A). When starved in the presence of PMSF, *sec3-2* cells showed accumulation of GFP-Atg8 puncta inside the vacuole at permissive temperature, but were present outside the vacuole at non-permissive temperature, suggesting an autophagy block prior to fusion with vacuole (Fig. S1E). In accordance with our GFP-Atg8 processing assay, *sec15-1* did not show any significant accumulation of multiple Atg8 puncta (Fig. 2B).

**Figure 2:**
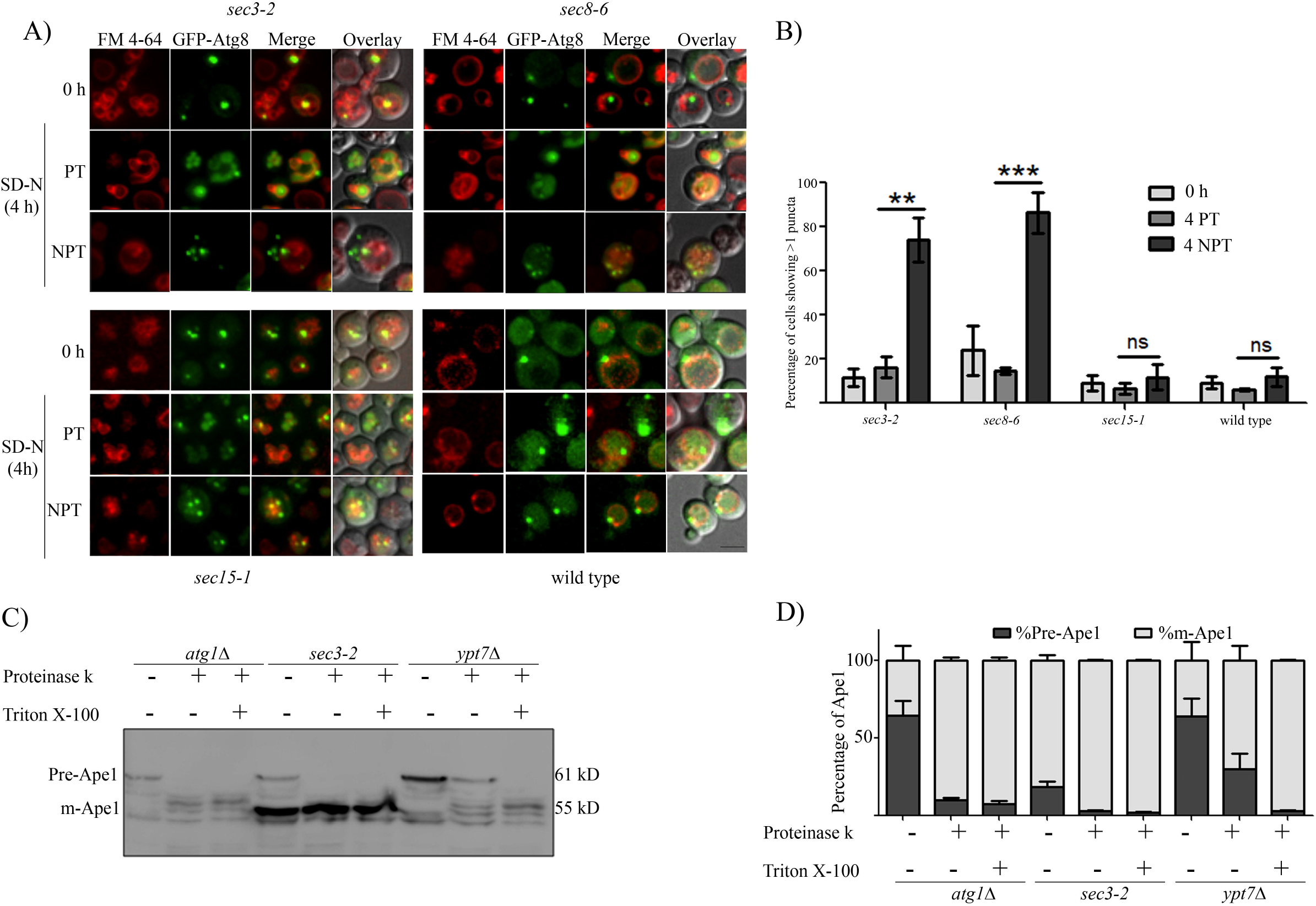
Mutants of exocyst complex accumulate incomplete autophagosomes. A) Cells expressing GFP-Atg8 were cultured in SD-Ura with FM 4-64 and moved to starvation at PT and NPT. Fluorescence microscopy images were acquired at 0hour and 4 hours in SD-N. Maximum intensity projected images are shown. Scale bar: 2μm. B) Cells showing more than one puncta in A were scored and means of three independent experiments are represented in the bar graph. A minimum of 100 cells were counted per experiment. Error bars represent standard error. Statistical significance was analysed by Student’s unpaired t-test. ns-nonsignificant, **, *P*<0.01, ***, *P*<0.001. C) *atg1*Δ, *ypt7*Δ and *sec3-2*cells were starved at NPT for 4 hours. Cells were harvested, spheroplasts were made and lysed. The clarified lysates were treated with either Proteinase K or Proteinase K with Triton X-100and analysed by immunoblotting using anti-Ape1 antibody. D) Intensities of precursor (Pr-Ape1) and mature Ape1 (m-Ape1) bands in each lane of C were measured using ImageJ (NIH). The percentages of Pr-Ape1 andm-Ape1 were determined and mean of three independent experiments plotted as bar graphs. Error bars represent S.E.M.

We wanted to test if the GFP-Atg8 puncta represent complete or incompletely formed autophagosomes. For this, we employed the protease protection assay using Ape1 maturation as a readout. Ape1 gets delivered as precursor form to the vacuole by Cvt (cytoplasm to vacuole targeting) vesicles in growth conditions and by autophagosomes during starvation, where it gets cleaved and converted into matureApe1. If the precursor form of Ape1 is entrapped in vesicles as in the case of *ypt*7Δ cells, it is resistant to the action of externally added protease such as Proteinase K. Addition of detergent, Triton X-100 dissolves the membrane of vesicles and exposes the precursor Ape1 to Proteinase K. However, if the autophagosomes are not formed as in *atg1*Δ cells or partially formed, Proteinase K can access and cleave precursor Ape1. We asked whether the autophagosomal structures (multiple GFP-Atg8 puncta) observed at non-permissive temperature in *sec3-2* cells were completely formed.

Similar to *atg1*Δ, *sec3-2* showed the presence of pre-Ape1 in untreated fraction but not in the fraction treated with either Proteinase K alone or Proteinase K with Triton X-100 (Fig. 2C). This data shows that at non-permissive temperature, the multiple autophagosomal structures in *sec3-2* were incompletely formed and therefore are not able to fuse with the vacuole.

We then investigated if the exocyst component Sec3 associates with autophagy components. Live cell microscopy revealed a dynamic interaction of Sec3-GFP and 2xmCherry-Atg8 (Fig. S2A, Video 1) and also with 2xmCherry-Atg9 (Fig. S2B).

### Atg9 vesicle trafficking is perturbed in exocyst temperature sensitive mutants

As we observed that the elongation or completion of autophagosomes was impaired in *sec3-2* mutant of exocyst, we investigated if dysfunctional Atg9 trafficking attributed to these defects in autophagosome formation. Atg9 is involved in contributing membrane to the growing phagophore aiding in autophagosome elongation and completion. To and fro trafficking of Atg9 vesicles takes place from peripheral membrane sources such as mitochondria, ER and Golgi to the PAS, delivering membrane for autophagosome formation. This shuttling of Atg9 is known to be critical for autophagy progression (36).

Wild type, *atg1*Δ and *sec3-2* cells expressing Atg9-GFP were imaged in growth medium and imaged again 2 hours post starvation. As the Atg9 vesicles are constantly moving to PAS (anterograde trafficking) and recycling back to cytoplasm (retrograde trafficking) wild type cells show a distribution of Atg9 puncta at peripheral membrane sources in the cytoplasm and at the PAS (5, 37, 38).In agreement with these reports, we observed a uniform distribution of Atg9 puncta in wild type cells in nutrient rich medium, under basal autophagy conditions, suggesting dynamic Atg9 trafficking (0 hour, Fig. 3A). Interestingly, a small but significant number of *sec3-2* cells showed single punctum of Atg9 indicating that the Atg9 trafficking dynamics may be perturbed (Fig. 3B). However, the Atg9 trafficking was completely restored when these mutant cells were transferred to starvation conditions where autophagy is induced (Fig.3B). It is known that Atg9 vesicles show an increased mobility during starvation conditions (11) and as a result the mild effect seen during basal autophagy may have been overridden during such autophagy inducing conditions in both permissive and non-permissive conditions. In addition to showing a prominent Atg9 punctum, *sec3-2* cells also display several puncta at non-permissive temperatures. In order to do a more precise analysis, we resorted to studying Atg9 trafficking using the TAKA (Transport of Atg9 in Knockout of Atg1) assay. The retrograde trafficking of Atg9 is dependent on the Atg1 complex and thus in *atg1*Δ cells, Atg9 vesicles reach but get arrested at PAS (WT vs *atg1*Δ, Fig.3A). We observed Atg9 trafficking in *sec3-2* cells where ATG1 has been deleted by shifting these cells to PT and NPT conditions in starvation medium. We noticed a significant reduction in number of *sec3-2 atg1*Δ cells showing single puncta at non-permissive temperature as compared to the permissive temperature implying defects in anterograde trafficking. These results indicate that the Atg9 vesicles are being driven to the PAS albeit at a reduced rate (Fig. 3C).

**Figure 3:**
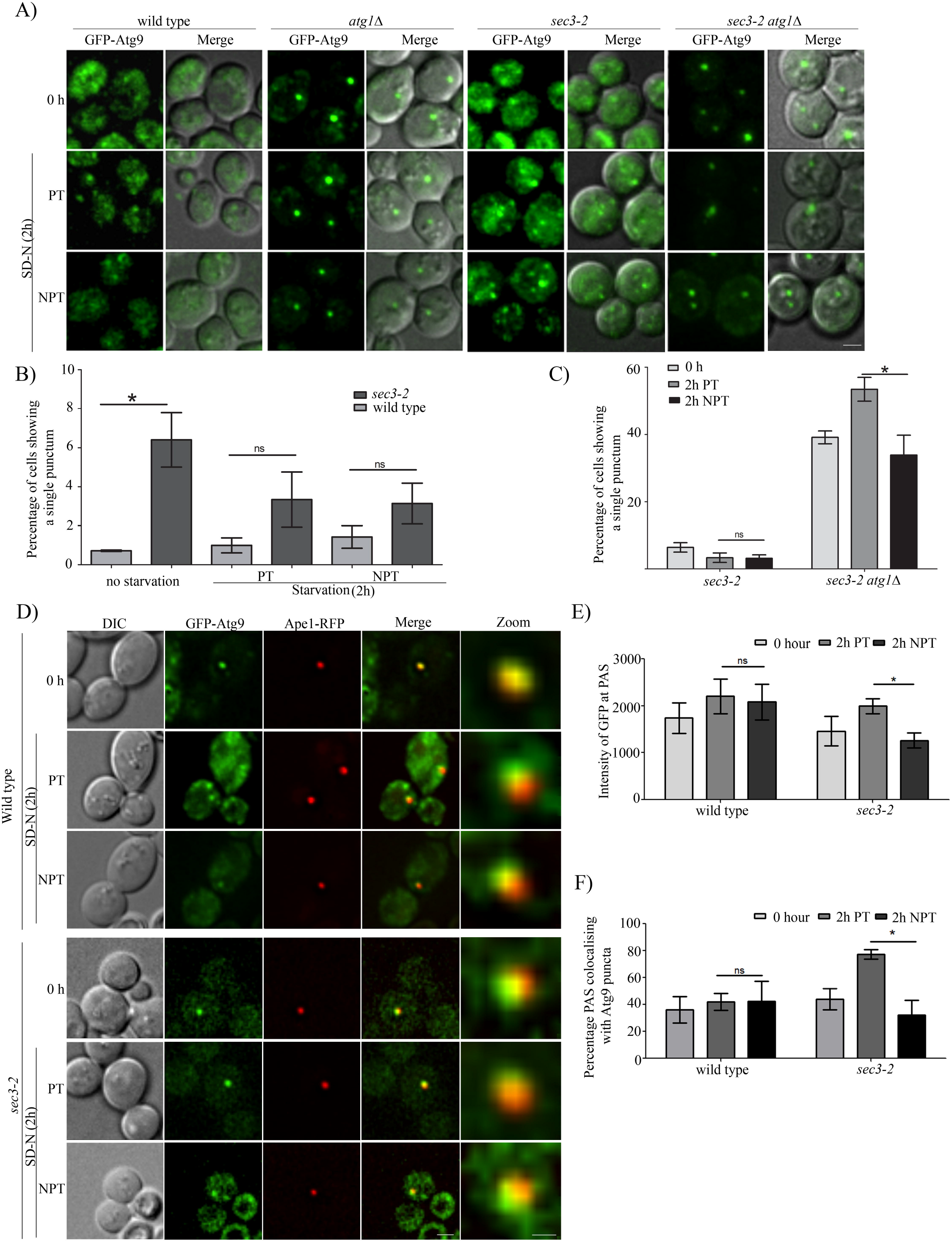
Anterograde trafficking of Atg9 vesicles is affected in an exocyst mutant. A) Wild type, *atg1*Δ,*sec3-2* and *sec3-2 atg1*Δ cells expressing GFP-Atg9 were grown in SD-Ura and then starved at PT or NPT. Fluorescence imaging was carried out at 0 hour and 2 hours. Deconvolved and maximum intensity projection images are shown. Scale bar: 2μm. B) Wild type and *sec3-2* cells as in A were scored for presence of single bright punctum of GFP-Atg9at 0 hour and 2 hours in starvation at PT and NPT. C) Comparison of *sec3-2* and *sec3-2 atg1*Δ cells showing single punctum of GFP-Atg9 as in A. For B and C more than 100 cells per three independent experiments were manually scored and mean values plotted with standard error. D) Wild type and*sec3-2*cells expressing GFP-Atg9 and Ape1-RFP were starved at PT or NPT. Fluorescence images were captured at 0 and 2 hours in SD-N. Scale bar: 2 μm (merge) and 0.5 μm (zoom). Intensity of GFP-Atg9 at PAS (marked by Ape1-RFP) was measured and average intensity of GFP-Atg9 is plotted in E. F) The percentage of PAS that colocalize with bright Atg9 puncta was determined from D and mean values were plotted. Error bars represent S.E.M. Statistical significance was analysed by Student’s unpaired t-test. ns-nonsignificant, *, *P*<0.05.

To further confirm these observations, we measured the amount of Atg9 at PAS as a read-out of Atg9 trafficking flux. We reasoned that as the retrograde trafficking was unaffected in the exocyst mutant, any changes in Atg9 levels at PAS would reflect a change in anterograde Atg9 dynamics. We co-expressed GFP-Atg9 and Ape1-RFP which served as a marker for PAS. We imaged cells at 2 hours post starvation in permissive and non-permissive conditions (Fig. 3D). The images with colocalised GFP-Atg9 and Ape1-RFP puncta were then used for intensity measurements. In agreement with our earlier results, we found that the average intensity of Atg9 at PAS was indeed reduced in *sec3-2* but not in WT cells (NPT vs PT, Fig. 3E). Furthermore, this mutant showed decreased occurrence of Atg9 at the PAS (PT vs NPT, Fig. 3F). These results indicate that in addition to an ineffective anterograde transport of Atg9 vesicles from peripheral sources to PAS, a dysfunctional exocyst component also led to a decreased number of Atg9 vesicles being tethered at PAS.

### The autophagy specific exocyst complex is distinct from its secretory pathway counterpart

As described previously, we noticed that mutants of some of the exocyst subunits did not show any effect on autophagy (Fig. 1D). This observation led us to consider the possibility of a disparity in the autophagy function associated exocyst complex than what is known for the secretory associated exocyst complex. To test this hypothesis, we resorted to size exclusion chromatography to analyse the exocyst complex size under starvation conditions.

For our initial analysis, we subjected Sec8-GFP cell lysates collected from cells grown in nutrient rich or starvation medium to size exclusion chromatography followed by western blotting with anti-GFP antibody. Quantification of intensity on western blot yielded a single peak from cells cultured in rich medium while those from starvation medium showed two distinct peaks (Fig. 4 A and 4B). To verify these results, we repeated the experiment with other subunits of the exocyst complex in starvation conditions. Interestingly, we find that the lysates of Sec5-GFP, Sec6-GFP and Sec10-GFP showed the presence of two peaks as seen for Sec8-GFP (Fig. 4B, starvation). However, Sec15-GFP, Exo70-GFP and Exo84-GFP showed only the first peak as seen for Sec8-GFP lysates from growth conditions (Fig.4C and 4D). These results suggest that the first peak corresponds to the peak seen in growth medium, while the second one suggested the presence of a smaller complex that was unique to autophagy inducing conditions. Importantly, only the exocyst subunits that were affecting autophagy showed the second peak.

**Figure 4:**
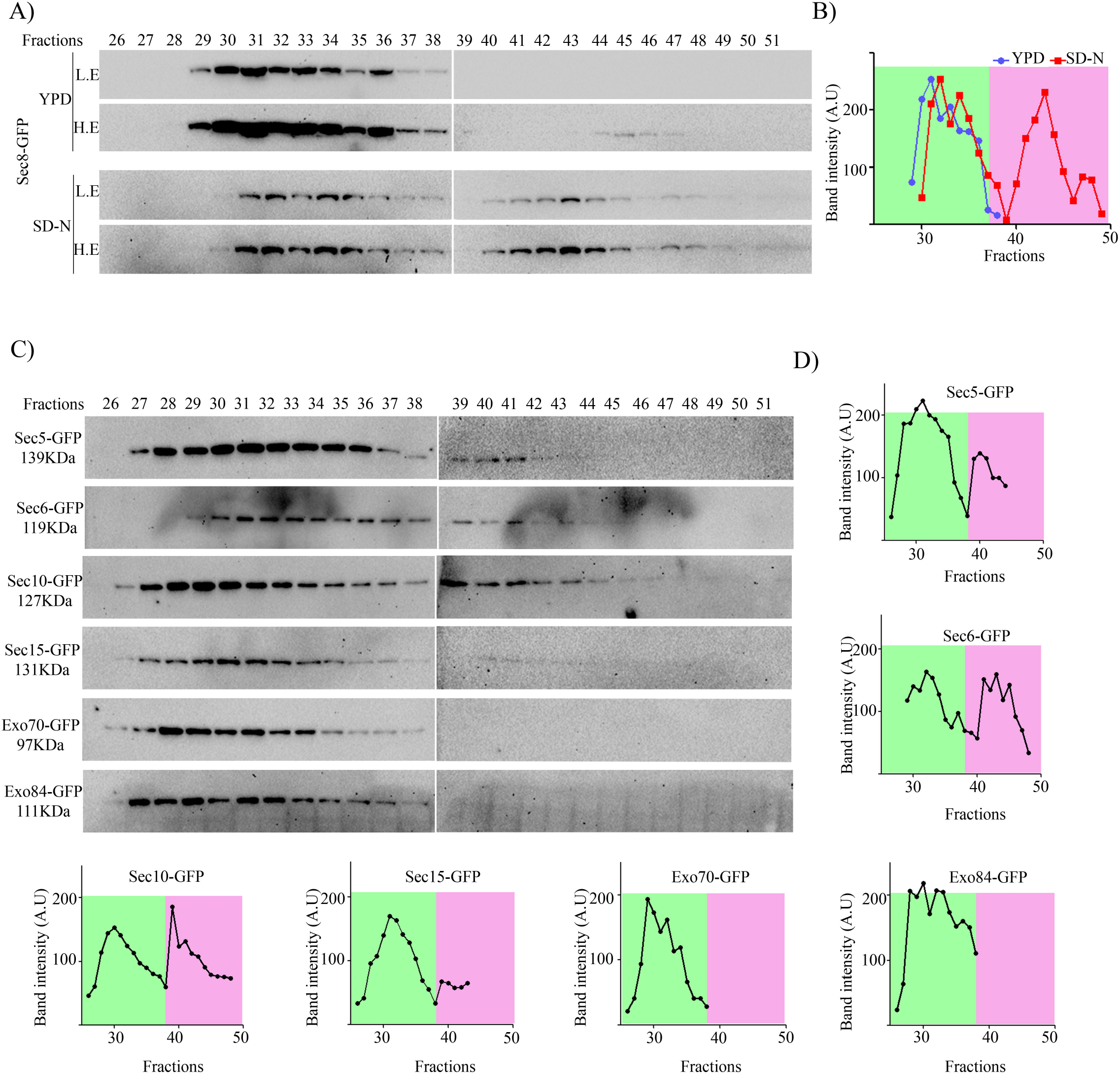
Autophagy prevalent conditions reveal presence of a subcomplex of exocyst comprising of subunits that are required for autophagy. A) Sec8-GFP cells were grown in rich medium (YPD) and starved for 4 hours. Clarified supernatant (cytosol) from YPD or SD-N grown cells were prepared and subjected to size exclusion chromatography using Superose 6 high load 10/300GL column. Fractions were collected and analysed by western blotting using anti-GFP antibody. L.E-lower exposures and H.E-higher exposures. B) Intensities of bands from A were quantitated and plotted against fractions. Peak in green area represent higher molecular weight exocyst complex associated with secretory function while peak in pink area represent starvation specific exocyst subcomplex. C) Strains expressing exocyst subunits tagged with GFP were starved for 4 hours and processed as in A. Fractions were analysed by western blotting. Intensity of bands from these western blots is plotted in D.

## Discussion

Autophagy is a complex catabolic process involving formation of double membrane vesicles-autophagosomes. The biogenesis of autophagosomes involve several membrane related transport events in which tethering complexes play an important role. We identified one such tethering complex called the exocyst to be involved in autophagy. The multisubunit exocyst complex has been characterized for its role in the secretory pathway. The secretory exocyst is an octameric complex and has a function of tethering post-Golgi secretory vesicles to the target membrane (39). Our study reveals the role of a subset of the exocyst complex proteins in the formation of autophagosomes. We systematically address the role of the exocyst subunit proteins for their subunit specific involvement and their potential role in autophagy.

In our study, we show that not all the subunits of exocyst complex are indispensable for its function in autophagy. Our experiments with temperature sensitive (Ts) strains of *S. cerevisiae* revealed that the mutants of several subunits of exocyst complex (Sec3, Sec5, Sec6, Sec8 and Sec10), but not others (Sec15, Exo70 and Exo84), showed defect in selective and general autophagy. Further investigation by fluorescence microscopy of these mutants showed presence of more than one GFP-Atg8 puncta in cells at non-permissive temperature indicating that the autophagosomes were possibly formed. These puncta accumulated outside the vacuole and therefore confirmed our observation that the autophagy block in these mutants was in a step prior to fusion with the vacuole. Interestingly, these Atg8 puncta represented incompletely formed autophagosomes as revealed by the protease protection assay suggesting a potential role for exocyst in autophagosome expansion and completion as well.

One of the key requirements for autophagosome expansion is delivery of membrane by Atg9 vesicles to the site of autophagosome biogenesis (PAS) (5). Our investigation in this direction revealed several interesting defects in the trafficking of Atg9 from the peripheral membrane sources to the PAS in the exocyst mutant. Firstly, we observed less colocalization events between Atg9 and PAS. Secondly, among these colocalised events, there was decreased occupancy of Atg9 vesicles at PAS. Thirdly, although retrograde trafficking appeared unperturbed, anterograde movement of Atg9 vesicles to PAS from peripheral sources appeared to be compromised, explaining the reason for decreased presence of Atg9 at the PAS. Put together, these observations point at impaired Atg9 trafficking from peripheral membrane sources to the PAS. Thus, this decreased rate of delivery of Atg9 vesicles but not their recycling/exit from PAS, must mean that the membrane supply to the phagophore is limited. Restricted Atg9 trafficking has been reported to clamp down on autophagosome biogenesis (11). We propose that diminished supply of Atg9 membrane sources may result in abortive autophagosome biogenesis in these mutants. This limited supply may allow initiation of autophagosome biogenesis but may not be sufficient to fuel expansion and completion of autophagosomes. This may explain the presence of multiple incompletely formed autophagosomes in these mutants.

The next question we addressed was why only certain members of this complex affected autophagy. A previous report suggested that in mammalian cells two subcomplexes of exocyst, a Sec5 containing complex that suppresses autophagy, and an autophagy activating complex consisting of Exo84 exist (32). RalB-a Ral GTPase, promotes autophagy induction by triggering a switch in the association of autophagy initiation complex (ULK1/Beclin1/VPS34 complex) from the Sec5 subcomplex of exocyst to Exo84 subcomplex. The presence of such distinct subcomplexes has also been reported in yeast, wherein during secretion, exocyst exists as two subcomplexes-one for targeting and other for bridging secretory vesicles with targeting complex (31). The assembly of complete complex is regulated by the Sec4 and Rho1 GTPases (27, 29, 40). Our biochemical investigations using size exclusion chromatography also indicates the possibility of an autophagy specific exocyst complex that differs in composition from that of the conventional secretory exocyst complex components. These results are in agreement with our genetic mutant based analysis which suggest a disparity in requirement of the exocyst complex subunits for autophagy function. Absence of functional Sec15, Exo70 and Exo84 did not affect autophagy flux progression. Interestingly, the same three subunits were found to be absent from the autophagy specific exocyst complex. Combined together, these results show that the constituents of exocyst complex functioning in autophagy is different from the one present in secretory pathway and harbours lesser components (5 out of 8). It is possible that the autophagy specific complex may have additional components.

In conclusion, our study highlights the moonlighting functions of some members of the exocyst machinery that form an autophagy specific subcomplex that catalyses membrane supply for autophagosome biogenesis.

## Materials and Methods

### Yeast strains and media

The detailed list of Strains used in this study is provided in Table 1.

**Table 1:**
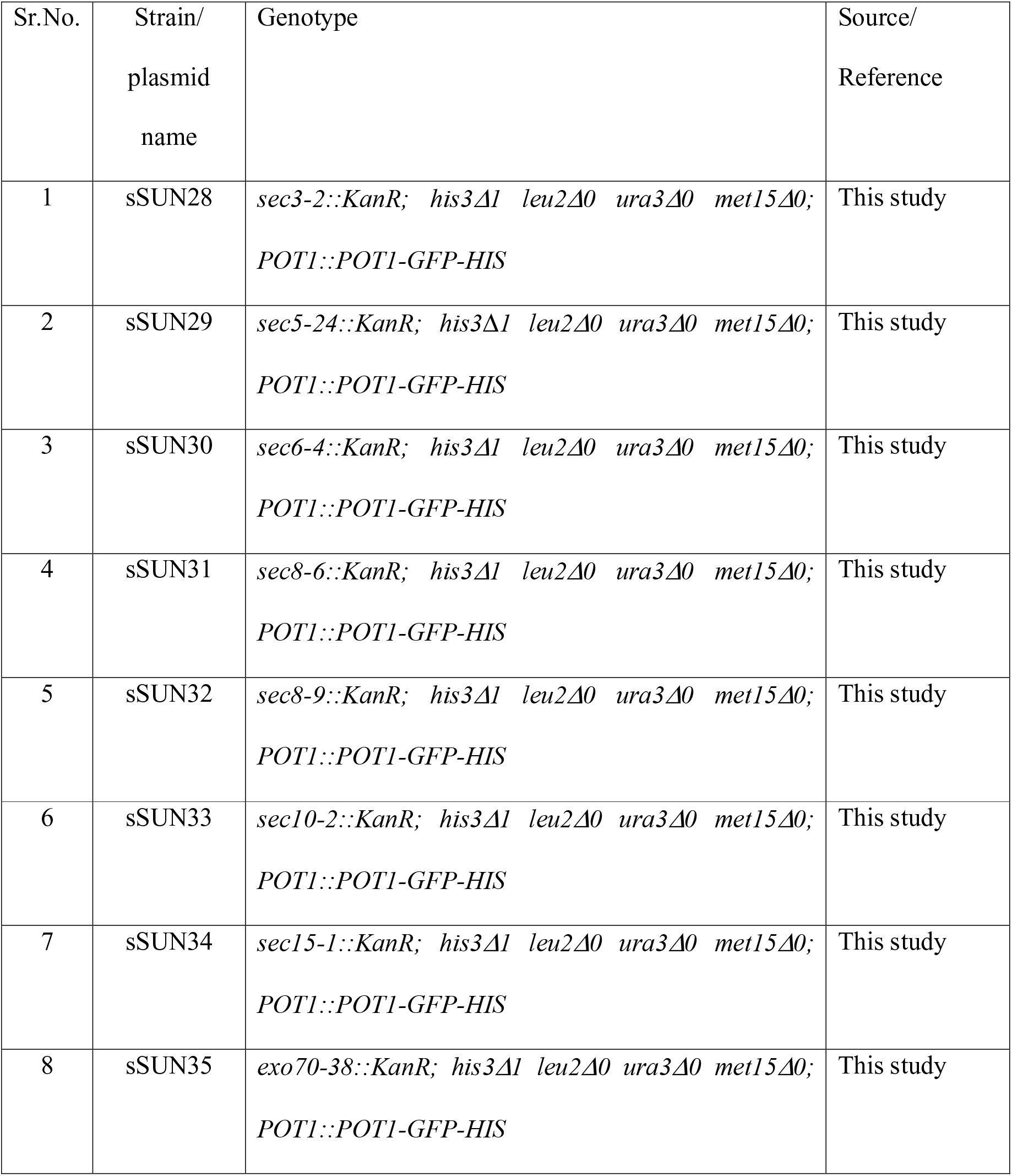
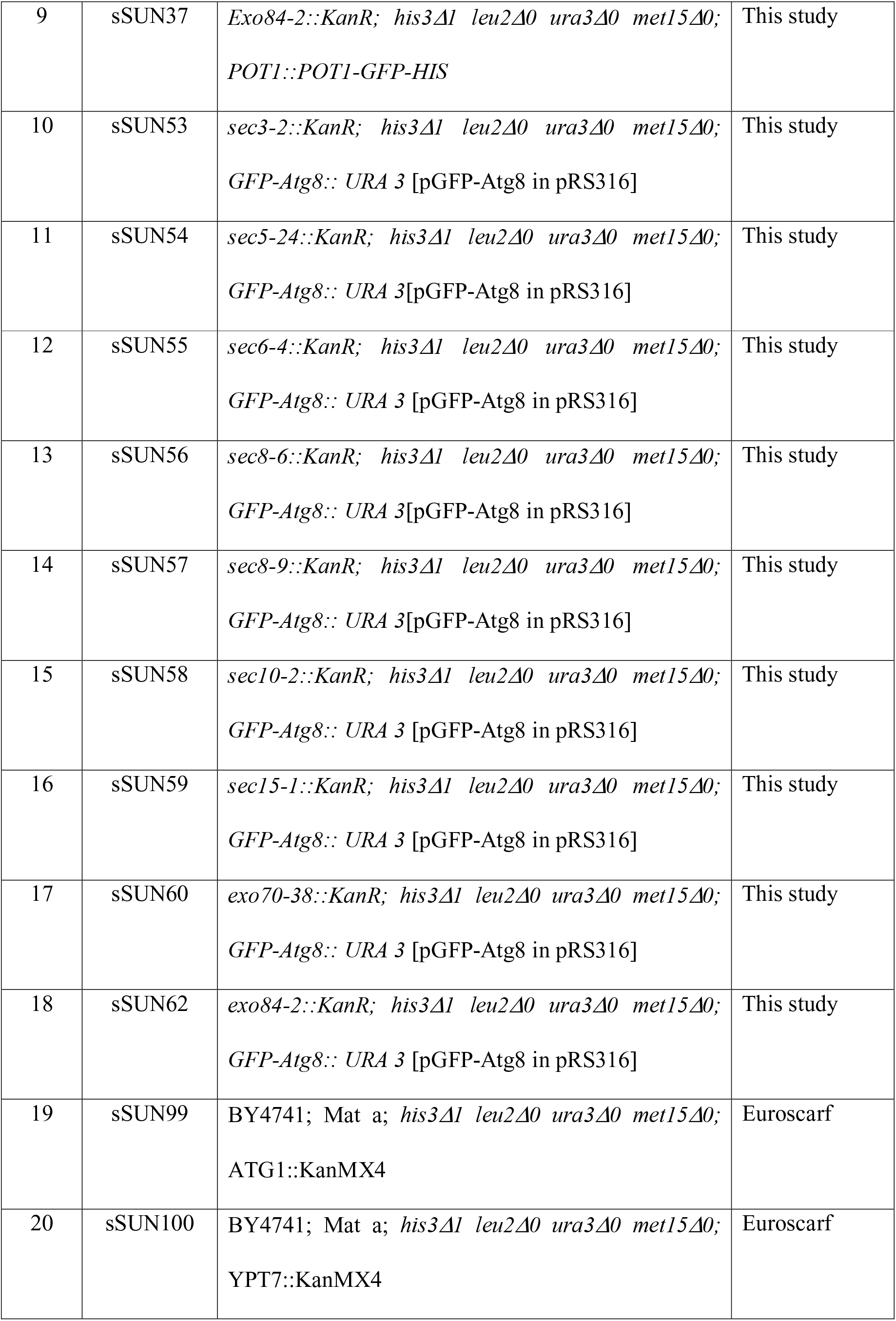
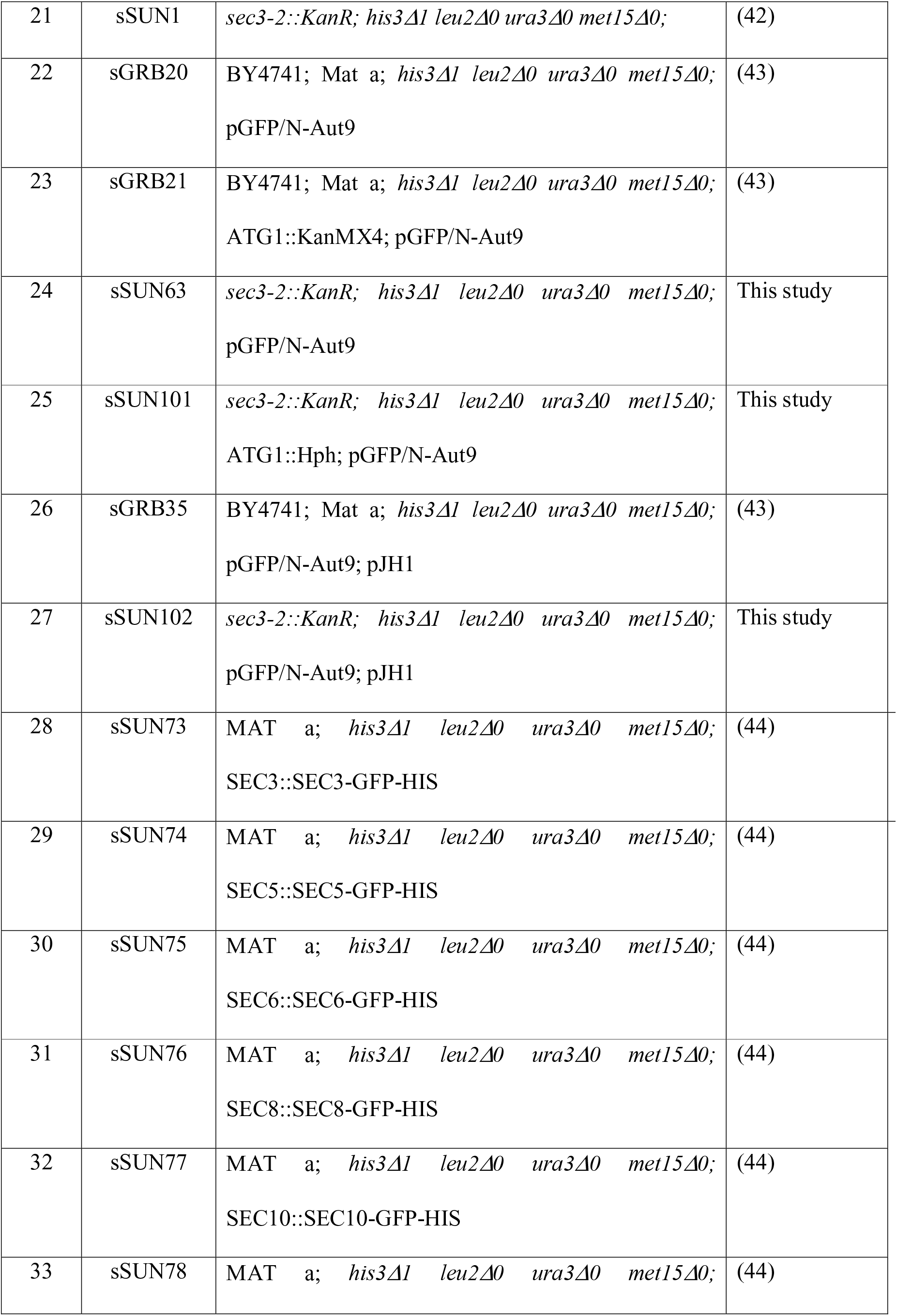
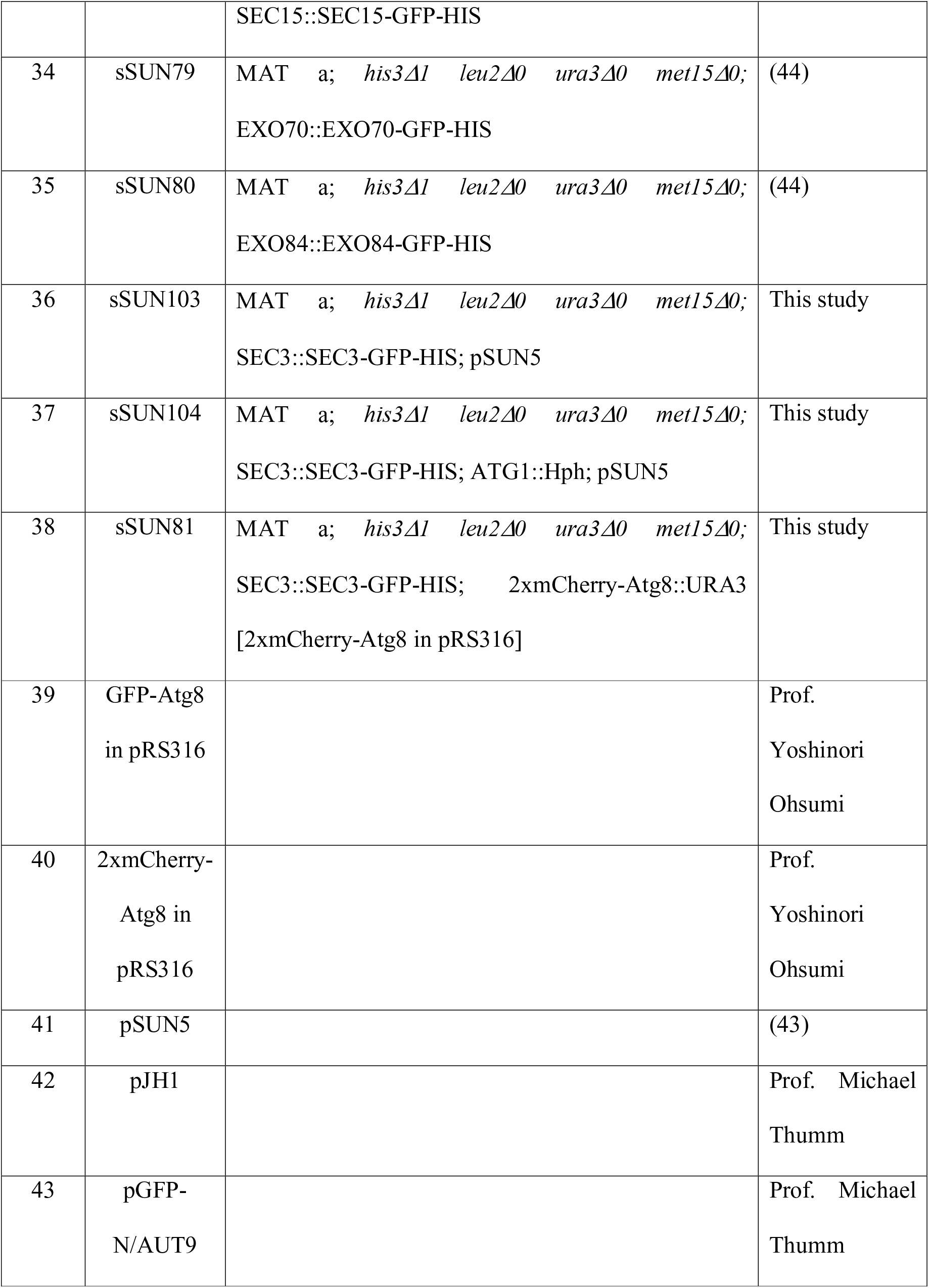
List of strains and plasmids used in this study:

Yeast cells were grown in YPD media (1% yeast extract, 2% peptone and 2% dextrose) or selection media SD-X (0.17% yeast nitrogenous base, 0.5% ammonium sulphate, 2% dextrose, 0.002% uracil, 0.02% histidine, 0.02% methionine, 0.015%lysine and 0.01% leucine), X being the amino acid to be exempted. Cells were starved by incubation in starvation media (SD-N) (0.17% yeast nitrogenous base and 2% dextrose). Oleate medium (2.64mM K_2_HPO_4_, 17.36mM KH_2_PO_4_, pH6.0, 0.1% oleic acid with 0.5% Tween-40, 0.25% yeast extract and 0.5% peptone) was used to induce peroxisomes. Wild type and knockout strains were transformed using Lithium acetate method (41). For temperature sensitive strains the cells were incubated at 25^□^C overnight in transformation mix instead of being subjected to a heat shock at 40^□^C.

All wild type and knockout strains were grown at 30^□^C while temperature sensitive strains were grown at 25^□^C. Assays were performed at permissive temperature (25^□^C) or non-permissive temperature (37^□^C).

### Pexophagy Assay

Wild type yeast cells expressing Pot1-GFP were a kind gift from Prof. Rachubinski, University of Alberta, Canada. Temperature sensitive strains were transformed with Pot1-GFP-HIS cassette which was PCR amplified from the genomic DNA of WT POT1-GFP cells. Actively growing cells expressing Pot1-GFP were transferred to oleate medium (A_600_=1) to induce peroxisome biogenesis (for microscopy, FM4-64 (1μg/ml) was added to label the vacuoles). These cells were washed twice with water and starved in SD-N medium (A_600_=3). Cells were collected at mentioned time intervals and either processed for western blotting or fluorescence microscopy.

### GFP-Atg8 processing assay

Cells were transformed with the plasmid GFP-Atg8 cloned in pRS316 vector backbone (a kind gift from Prof. Yoshinori Ohsumi). All strains expressing GFP-Atg8 were grown in SD-Ura medium and actively growing cells were transferred to starvation medium (SD-N). Cells were harvested at indicated time intervals and TCA precipitated. These precipitates were subjected to SDS-PAGE followed by western blotting.

### Fluorescence microscopy

Cells were mounted on 2% agarose pads on slides. Imaging was performed on DeltaVision microscope (GE Healthcare) using 60X or 100X objective and images were captured with the cool-SNAP HQ camera. Deconvolution and intensity projection of images was done using softWoRx software (GE Healthcare). Co-localisation analysis was performed manually or by softWoRx software. Brightness contrast adjustments were made for representation purpose in figures. Quantitation of number of cells showing a particular phenotype or intensity measurements were done using Fiji software (NIH).

### Protease Protection Assay

Cells A_600_= 60 were transferred to starvation medium and incubated at 37^□^for 4 hours. Cells were then harvested, washed once and resuspended in 30ml wash buffer with β-ME (0.6μl/ml) and incubated for 10 minutes at 37^□^C, 180rpm. Cells were then resuspended in spheroplasting buffer (SP) with zymolyase-20T (20mg/ml) and incubated at 37^□^C for ~1hour at 70 rpm. The resulting spheroplasts were washed once with SP buffer and lysed by incubating on ice for 5 minutes. The lysate was centrifuged at 300*g* for 10 minutes at 4°C. The supernatant was divided into 3 tubes for control, Proteinase K (40 μg/ml) and Proteinase K with Triton X-100 (0.2%) treatments. The reactions were incubated on ice for 15 minutes and then stopped by addition of TCA to a final concentration of 10% and samples were frozen at -80^□^C. The TCA precipitates were washed with acetone, air dried and resuspended in 25μl SDS-PAGE dye. Samples were separated on 8% SDS-PAGE gels followed by western blotting.

### Western blotting

TCA precipitates were washed with 80% acetone, air dried and boiled with SDS-PAGE dye. A_600_= 0.6 culture equivalents were loaded onto SDS-PAGE and transferred to PVDF membranes at 100V for 1-1.5 hours. Mouse anti-GFP monoclonal antibody at 1:3000 (Roche Applied Sciences), anti-Ape1 1:5000 (a kind gift from Prof. Yoshinori Ohsumi), anti-mouse and anti-rabbit secondary antibodies conjugated to HRP (Bio-Rad) were used. Blots were developed in G-box Chemi XT4 (Syngene) and band intensities were analysed using Fiji software (NIH).

### Gel filtration

Cultures grown in 2 litres YPD to A_600_= 0.8-1 were harvested or washed and starved in SD-N at A_600_=3 for 4 hours before harvesting. Cell pellets were then resuspended in lysis buffer (20mM HEPES pH 7.4, 100mM NaCl, 1 mM EDTA, 10mM β-mercaptoethanol, 0.5% NP-40, protease inhibitor cocktail). Lysis was done by vortexing the cell suspension along with 0.5mm glass beads (Sigma Aldrich) for 1 minute followed by 2 minutes incubation on ice, for 8-10 times. Lysate was then centrifuged at 13000*g* at 4^□^C for 10 minutes and the resultant supernatant was concentrated before loading onto equilibrated (20mM HEPES pH 7.4, 100mM NaCl) Superose 6 high load 10/300 column (GE Healthcare). The flow rate was kept constant at 0.2 ml/ minute. Fractions (250μl) were collected and TCA precipitated before loading on 8% SDS-PAGE gels. Western blotting was done and the relative band intensities were calculated using Fiji software and intensity values were plotted using GraphPad Prism software.

### Statistical Analysis and Image preparation

All statistical analysis was performed by using GraphPad Prism (GraphPad Software). Mean of three independent experiments were compared using unpaired two-tailed Student’s t-test. Images were processed post acquisition and prepared using softWoRx software (GE Healthcare). Brightness contrast adjustments were done for visualisation purpose during image preparation. Images were then collated using Adobe Photoshop CC.

---

**Figure S1: *sec8-9* mutant shows a block in pexophagy and general autophagy**

A) Peroxisome biogenesis was induced in *sec8-9*Pot1-GFP cells by growing in oleate medium. Cells were then starved at PT or NPT, samples collected at times indicated and analysed by immunoblotting.

B) Pot1-GFP expressing *sec8-9*cells were imaged at 2 hours in SD-N at PT or NPT. Green dots represent peroxisomes and red structures denote vacuoles labelled with FM 4-64. Images were deconvolved and projected for maximum intensity. Scale bar: 2μm.

C) Cells showing diffuse GFP vacuoles in B were manually scored. Bar graph represents the mean of three experiments with S.E.M.

D) GFP-Atg8 processing assay (similar to Fig.1D) was performed for *sec8-9* cells and analysed by immunoblotting.

E) *sec3-2* cells expressing GFP-Atg8 were starved in the presence of PMSF at PT and NPT for 4 hours and fluorescence images were acquired. Maximum intensity projection is shown. Scale bar: 2μm

**Figure S2: Colocalization of Sec3-GFP with 2xmCherry-Atg8 or2xmCherry-Atg9**

A) Cells expressing Sec3-GFP and 2xmCherry-Atg8 were grown in SD-URA medium and transferred to SD-N for 30 minutes. Cells were embedded on 2% agarose pads made in SD-N and imaged. Shown here are the snapshots of the supplementary video 1 at indicated time of appearance in the video. White arrows indicate partial colocalization and yellow arrows indicate complete colocalization. Scale bar: 2μm

B) Wild type and *atg1*Δ cells expressing Sec3-GFP and 2xmCherry-Atg9 were imaged at 0, 1 and 2 hours post starvation. Scale bar: 2μm and 0.5μm (Zoom). About 100 cells of wild type and *atg1*Δ in each experiment were manually scored for colocalization of Sec3-GFP and 2xmCherry-Atg9. Mean of percentage cells showing colocalization, as in C, from three independent experiments is plotted. Error bars represent S.E.M.

**Supplementary video 1: Colocalization of Sec3-GFP and 2xmCherry-Atg8**

Actively growing wild type cells expressing Sec3-GFP and 2xmCherry-Atg8 were transferred to starvation media (SD-N) and collected after 30 minutes. Live cell fluorescence imaging performed capturing 3 z sections. Shown in the video is single z section with an interval of 5 seconds across each frame. Scale bar: 2μm.

## Acknowledgements

We thank Prof. Charlie Boone (University of Toronto), Prof. Yoshinori Ohsumi (Tokyo Institute of Technology), Prof. Michael Thumm (Georg-August-Universität Göttingen), Dr. Kausik Chakraborty (Institute of Genomics and Integrative Biology), and Dr. Richard Rachubinski (University of Alberta) for sharing yeast strains, plasmids and antibodies. Thanks to Hariharan and Prem Anand Murugan (IIT-Kanpur) for the technical help in gel filtration experiments. We thank Aparna Hebbar, Vikas Yadav, Piyush Mishra and Gaurav Barve for critical reading of this manuscript and advice. This work was supported by Wellcome Trust/DBT India Alliance Intermediate Fellowship (509159-Z-09-Z) and intramural funds from JNCASR to RM. The authors declare no competing financial interests.

## Author contributions

RM conceived the idea. Most of the experiments were performed by SS with help from AA and SC. SS and RM analysed the data and wrote the manuscript. SM provided guidance for gel filtration experiments.

## References

1. Klionsky DJ, Emr SD. Autophagy as a regulated pathway of cellular degradation. Science. 2000;290(5497): 1717–21.

2. Zhi X, Feng W, Rong Y, Liu R. Anatomy of autophagy: from the beginning to the end. Cellular and molecular life sciences : CMLS. 2018;75(5):815–31.

3. Kim J, Huang NP, Stromhaug PE, Klionsky DJ. Convergence of multiple autophagy and cytoplasm to vacuole targeting components to a perivacuolar membrane compartment prior to de novo vesicle formation. The Journal of biological chemistry. 2002;277(1):763–73.

4. Suzuki K, Kirisako T, Kamada Y, Mizushima N, Noda T, Ohsumi Y. The pre-autophagosomal structure organized by concerted functions of APG genes is essential for autophagosome formation. The EMBO journal. 2001;20(21):5971–81.

5. Noda T, Kim J, Huang NP, Baba M, Tokunaga C, Ohsumi Y, et al. Apg9p/Cvt7p is an integral membrane protein required for transport vesicle formation in the Cvt and autophagy pathways. The Journal of cell biology. 2000;148(3):465-80.

6. Hailey DW, Rambold AS, Satpute-Krishnan P, Mitra K, Sougrat R, Kim PK, et al. Mitochondria supply membranes for autophagosome biogenesis during starvation. Cell. 2010;141(4):656–67.

7. Mari M, Griffith J, Rieter E, Krishnappa L, Klionsky DJ, Reggiori F. An Atg9-containing compartment that functions in the early steps of autophagosome biogenesis. The Journal of cell biology. 2010;190(6):1005–22.

8. Ohashi Y, Munro S. Membrane delivery to the yeast autophagosome from the Golgi-endosomal system. Molecular biology of the cell. 2010;21(22):3998–4008.

9. Ravikumar B, Moreau K, Jahreiss L, Puri C, Rubinsztein DC. Plasma membrane contributes to the formation of pre-autophagosomal structures. Nature cell biology. 2010;12(8):747–57.

10. Reggiori F, Shintani T, Nair U, Klionsky DJ. Atg9 cycles between mitochondria and the pre-autophagosomal structure in yeasts. Autophagy. 2005;1(2):101–9.

11. Yamamoto H, Kakuta S, Watanabe TM, Kitamura A, Sekito T, Kondo-Kakuta C, et al. Atg9 vesicles are an important membrane source during early steps of autophagosome formation. The Journal of cell biology. 2012;198(2):219–33.

12. Hamasaki M, Noda T, Ohsumi Y. The early secretory pathway contributes to autophagy in yeast. Cell structure and function. 2003;28(1):49–54.

13. Tan D, Cai Y, Wang J, Zhang J, Menon S, Chou HT, et al. The EM structure of the TRAPPI11 complex leads to the identification of a requirement for COPII vesicles on the macroautophagy pathway. Proceedings of the National Academy of Sciences of the United States of America. 2013; 110(48): 19432–7.

14. Yen WL, Shintani T, Nair U, Cao Y, Richardson BC, Li Z, et al. The conserved oligomeric Golgi complex is involved in double-membrane vesicle formation during autophagy. The Journal of cell biology. 2010; 188(1): 101–14.

15. Lemus L, Ribas JL, Sikorska N, Goder V. An ER-Localized SNARE Protein Is Exported in Specific COPII Vesicles for Autophagosome Biogenesis. Cell reports. 2016;14(7):1710–22.

16. Nair U, Jotwani A, Geng J, Gammoh N, Richerson D, Yen WL, et al. SNARE proteins are required for macroautophagy. Cell. 2011;146(2):290–302.

17. Zhao S, Li CM, Luo XM, Siu GK, Gan WJ; Zhang L, et al. Mammalian TRAPPIII Complex positively modulates the recruitment of Sec13/31 onto COPII vesicles. Scientific reports. 2017;7:43207.

18. Brunet S, Shahrzad N, Saint-Dic D, Dutczak H, Sacher M. A trs20 mutation that mimics an SEDT-causing mutation blocks selective and non-selective autophagy: a model for TRAPP III organization. Traffic. 2013;14(10):1091–104.

19. Lipatova Z, Majumdar U, Segev N. Trs33-Containing TRAPP IV: A Novel Autophagy-Specific Ypt1 GEF. Genetics. 2016;204(3):1117–28.

20. Taussig D, Lipatova Z, Segev N. Trs20 is required for TRAPP III complex assembly at the PAS and its function in autophagy. Traffic. 2014;15(3):327–37.

21. Wang J, Menon S, Yamasaki A, Chou HT, Walz T, Jiang Y, et al. Yptl recruits the Atgl kinase to the preautophagosomal structure. Proceedings of the National Academy of Sciences of the United States of America. 2013;110(24):9800–5.

22. TerBush DR, Maurice T, Roth D, Novick P. The Exocyst is a multiprotein complex required for exocytosis in *Saccharomyces cerevisiae*. The EMBO journal. 1996;15(23):6483–94.

23. Pfeffer SR. Transport-vesicle targeting: tethers before SNAREs. Nature cell biology. 1999;1(1):E17–22.

24. Mei K, Li Y, Wang S, Shao G, Wang J, Ding Y, et al. Cryo-EM structure of the exocyst complex. Nature structural & molecular biology. 2018;25(2):139–46.

25. Munson M, Novick P. The exocyst defrocked, a framework of rods revealed. Nature structural & molecular biology. 2006;13(7):577–81.

26. Heider MR, Gu M, Duffy CM, Mirza AM, Marcotte LL, Walls AC, et al. Subunit connectivity, assembly determinants and architecture of the yeast exocyst complex. Nature structural & molecular biology. 2016;23(1):59–66.

27. Guo W, Sacher M, Barrowman J, Ferro-Novick S, Novick P. Protein complexes in transport vesicle targeting. Trends in cell biology. 2000;10(6):251–5.

28. Finger FP, Hughes TE, Novick P. Sec3p is a spatial landmark for polarized secretion in budding yeast. Cell. 1998;92(4):559–71.

29. Guo W, Roth D, Walch-Solimena C, Novick P. The exocyst is an effector for Sec4p, targeting secretory vesicles to sites of exocytosis. The EMBO journal. 1999;18(4):1071–80.

30. Luo G, Zhang J, Guo W. The role of Sec3p in secretory vesicle targeting and exocyst complex assembly. Molecular biology of the cell. 2014;25(23):3813–22.

31. Moskalenko S, Tong C, Rosse C; Mirey G; Formstecher E; Daviet L, et al. Ral GTPases regulate exocyst assembly through dual subunit interactions. The Journal of biological chemistry. 2003;278(51):51743–8.

32. Bodemann BO, Orvedahl A, Cheng T, Ram RR, Ou YH, Formstecher E, et al. RalB and the exocyst mediate the cellular starvation response by direct activation of autophagosome assembly. Cell. 2011; 144(2):253–67.

33. He B, Guo W. The exocyst complex in polarized exocytosis. Current opinion in cell biology. 2009;21(4):537–42.

34. Grote E, Carr CM, Novick PJ. Ordering the final events in yeast exocytosis. The Journal of cell biology. 2000;151(2):439–52.

35. Manjithaya R, Jain S, Farre JC, Subramani S. A yeast MAPK cascade regulates pexophagy but not other autophagy pathways. The Journal of cell biology. 2010;189(2):303–10.

36. Mari M, Reggiori F. Atg9 trafficking in the yeast *Saccharomyces cerevisiae*. Autophagy. 2007;3(2):145–8.

37. Lang T, Reiche S, Straub M, Bredschneider M, Thumm M. Autophagy and the cvt pathway both depend on AUT9. Journal of bacteriology. 2000;182(8):2125–33.

38. Young AR, Chan EY, Hu XW, Kochl R, Crawshaw SG, High S, et al. Starvation and ULK1-dependent cycling of mammalian Atg9 between the TGN and endosomes. Journal of cell science. 2006;119(Pt 18):3888–900.

39. Heider MR, Munson M. Exorcising the exocyst complex. Traffic. 2012;13(7):898–907.

40. Guo W, Grant A, Novick P. Exo84p is an exocyst protein essential for secretion. The Journal of biological chemistry. 1999;274(33):23558–64.

41. Gietz RD, Woods RA. Transformation of yeast by lithium acetate/single-stranded carrier DNA/polyethylene glycol method. Methods in enzymology. 2002;350:87–96.

42. Li Z, Vizeacoumar FJ, Bahr S, Li J, Warringer J, Vizeacoumar FS, et al. Systematic exploration of essential yeast gene function with temperature-sensitive mutants. Nature biotechnology. 2011;29(4):361–7.

43. Barve G, Sridhar S, Aher A, Sahani MH, Chinchwadkar S, Singh S, et al. Septins are involved at the early stages of macroautophagy in S. cerevisiae. Journal of cell science. 2018;131(4).

44. Huh WK, Falvo JV, Gerke LC, Carroll AS, Howson RW, Weissman JS, et al. Global analysis of protein localization in budding yeast. Nature. 2003;425(6959):686–91.

